# Beta spectral power during sleep is associated with impaired recall of extinguished fear

**DOI:** 10.1101/2022.09.08.506006

**Authors:** Dan Denis, Ryan Bottary, Tony J. Cunningham, Sean P.A. Drummond, Laura D. Straus

## Abstract

The failure to retain memory for extinguished fear plays a major role in the maintenance of post-traumatic stress disorder (PTSD), with successful extinction recall necessary for symptom reduction. Disturbed sleep, a hallmark symptom of PTSD, impairs fear extinction recall. However, our understanding of the electrophysiological mechanisms underpinning sleep’s role in extinction retention remain underdetermined. We examined the relationship between the microarchitecture of sleep and extinction recall in healthy humans (n=46, both male and females included) and a pilot study in individuals with PTSD (n=12). Participants underwent a fear conditioning and extinction protocol over two days, with sleep recording occurring between conditioning and extinction. Twenty-four hours after extinction learning, participants underwent extinction recall. Power spectral density (PSD) was computed for pre- and post-extinction learning sleep. Increased beta band PSD (∼17-26Hz) during pre-extinction learning sleep was associated with worse extinction recall in healthy participants (*r*=.41, *p*=.004). Beta PSD was highly stable across three nights of sleep (intraclass correlations (ICC)>0.92). Individuals with PTSD were found to have increased beta PSD compared to healthy controls (*p*s < .039), and beta PSD correlated with extinction recall in the PTSD group at a similar magnitude to controls (*r*=.39). Results suggest beta band PSD is elevated in PTSD, and is specifically implicated in difficulties recalling extinguished fear.

**Significance statement:** Disturbed sleep is a hallmark feature of posttraumatic stress disorder (PTSD). Certain neural oscillations that occur during sleep have been shown to be altered in PTSD. These include increased 15-30Hz beta oscillations, which are believed to index cortical hyperarousal. Alongside sleep disturbances, patients also show difficulty in recalling extinguished fear. Although prior studies have linked sleep with extinction retention, no studies have investigated the role that neural oscillations during sleep play in this process. Here, we show in both healthy participants and PTSD patients, that increased beta oscillatory power during sleep is associated with impaired extinction retention. Therefore, this study provides new evidence that electrophysiological changes in the sleep EEG of PTSD patients is also implicated in extinction recall processes.

## Introduction

Post-traumatic stress disorder (PTSD) is an emotional disorder characterized by a hyperarousal syndrome that impacts both daytime functioning and sleep ^1^. PTSD has been described as a disorder of emotional learning and memory that can be modeled experimentally using Pavlovian aversive conditioning paradigms ^2–4^. Such paradigms pair aversive stimuli with neutral stimuli, generating a measurable conditioned fear response. This learning process parallels the maladaptive fear learning in PTSD, whereby neutral situations, locations or objects become associated with an aversive outcome (i.e., the index trauma). Fearful responses to non-threatening stimuli can be overcome by a process of extinction learning, whereby individuals are repeatedly presented with conditioned stimuli in the absence of the aversive outcome. Memory for learned extinction attenuates fear responses ^4^ and is necessary for PTSD symptom reduction ^5^.

Sleep disturbances are considered a hallmark figure of PTSD ^6,7^, and a growing body of research has linked sleep to extinction memory processes. Both rapid eye movement (REM) and non-REM (NREM) sleep following extinction learning has been linked to the consolidation and retention of extinction memory ^8–14^. Additionally, total sleep deprivation *prior* to extinction learning can disrupt later recall of extinguished fear, suggesting that sleep prior to extinction learning may be important in preparing the brain for the consolidation of new learning ^15^.

Theta (∼4-8Hz) spectral power during REM sleep has been linked to emotional memory processing ^16^, and has been shown to be elevated in trauma-exposed individuals who were resilient to, compared to those diagnosed with, PTSD (Cowdin et al., 2014). Beta spectral power (∼16-30Hz) has also been shown to be elevated in PTSD compared to those without PTSD ^18–21^, and has been theorized as a marker of disturbed sleep and hyperarousal ^22–25^. Although these signatures are potentially important biomarkers of PTSD, results are not always consistent across studies ^26–28^.

These literatures provide converging evidence for a critical role of sleep in fear extinction retention. In PTSD, sleep disturbances may play a key role in the difficulty for these individuals to retain memories of extinguished fear and thus maintain disordered symptoms. Here, we examined whether sleep oscillatory activity that is altered in PTSD, such as theta and beta spectral power, impact the basic fear memory processes that are implicated in PTSD development, such as the retention of extinction memories.

Utilizing an existing dataset, we first examined associations between pre-extinction learning sleep spectral power and extinction recall in healthy controls ^15^. Given that total sleep deprivation prior to extinction learning impairs later recall, we reasoned that differences in sleep microarchitecture *during* sleep prior to extinction may also play a role in later retention. As a secondary analysis, we also examined associations with *post*-extinction learning sleep spectral power, on the basis of other work showing sleep macroarchitecture correlations ^8,11–14^. Utilizing a second pilot dataset ^29^ we tested whether similar associations were present in a sample of individuals with PTSD. Finally, if these measures of sleep oscillatory activity reflect diagnostic biomarkers of PTSD, it is critical that they are consistently represented in the sleep record. As such, we conducted exploratory analyses examining the night-to-night consistency in potentially problematic spectral power identified through the primary analyses.

We predicted that in healthy controls, greater theta-band spectral activity during REM sleep would be associated with better extinction recall, given theta’s putative role in emotional memory processing. Because sleep disruption via deprivation impairs extinction recall ^15^, and greater beta-band spectral power during sleep is a proxy of disturbed sleep ^30^, we expected greater beta-band spectral activity during sleep (both REM and NREM) to be associated with poorer subsequent recall of extinguished fear. In the pilot group of individuals with PTSD, we expected decreased REM theta and increased REM/NREM beta activity compared to the healthy controls. We also predicted that associations between spectral power and extinction recall would be of the same magnitude and direction as observed in the healthy controls.

## Study 1

### Materials and methods

This study was a reanalysis of a previously published dataset (Marshall et al., 2014; Straus et al., 2017). Please see Straus et al., 2017 for the full protocol details. The investigation was carried out in accordance with the Declaration of Helsinki. The University of California, San Diego’s Human Research Protections Program approved the study. Informed consent was obtained from all participants.

### Participants

Seventy-three healthy adults were enrolled in the study, and 71 participants with full datasets were included in the final analysis. Participants were healthy adults aged 18-39 years old with no current medical or psychiatric diagnosis, and exhibited a consistent blink-startle response at screening (over 75% discernable responses to twelve 105dB 40ms startle pulses).

### Procedure

Participants spent 4 consecutive days and nights in the sleep laboratory (**Figure 1A**). Following an adaptation night of sleep, participants underwent a fear potentiated startle procedure (see below) consisting of three sessions: fear acquisition (day 1); fear extinction learning (day 2); and fear extinction recall (day 3, the focus of the present analysis). All testing took place in the evening. Participants were randomized to either: 1) a Normal sleep condition consisting of a full night of sleep on all experimental nights (*n*=21); 2) 36 hours of total sleep deprivation after extinction learning (“Post-extinction deprivation”, *n* = 25) or 3) 36 hours of total sleep deprivation before extinction learning (“Pre-extinction deprivation”; *n* = 25). Sleep was monitored using polysomnography (see below).

**Figure 1.**
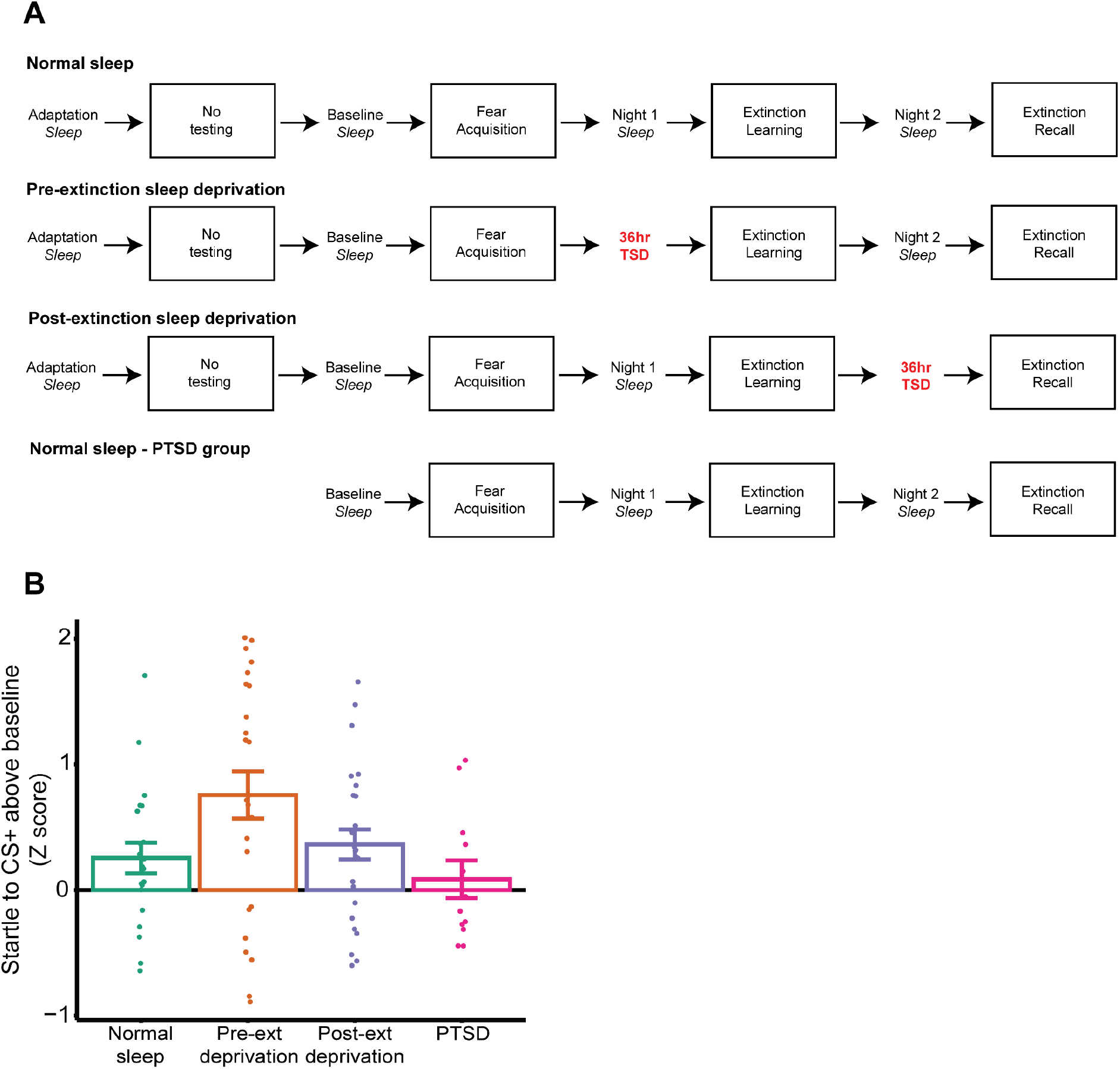
Experimental design and behavioral results. **A** - Study timeline for each group. **B** - Fear extinction recall, displayed as the startle response to CS+ above baseline (Z score). Note that the PTSD sample performed a different fear potentiated startle protocol. Error bars indicate the between-subject standard error.

### Fear potentiated startle

Each session began with six startle pulses (108db, 30ms acoustic startle probes) presented in the absence of any other stimuli in order for participants to acclimate startle responses. The fear acquisition session on day 1 consisted of 1) eight 6-second long presentations of a blue circle (CS+), following by a 500 ms electric shock (US) in 75% of blue circle trials; 2) eight 6-second presentations of a red circle serving as a second CS+, also followed by a 500 ms electric shock US in 75% of trials; 3) sixteen 6-second presentations of a yellow circle serving as a non-reinforced conditioned stimulus (CS-), never followed by a shock; and 4) sixteen presentations of the startle pulse in the absence of any stimuli (noise only; NA). The first half of the acquisition session consisted of only blue CS+ trials, and the second half consisted of only red CS+ trials. Startle pulses were presented 4 seconds following CS+ or CS-onset. Stimulus presentation was randomized within each CS+ acquisition block (blue vs red) and with the constraint of the two trials of each type (CS+, CS-, and NA) per block. On day 2, participants underwent extinction learning, consisting of 16 presentations of each stimulus type (blue CS+, yellow CS-, blank screen) in a block randomized order as in the acquisition session. No shocks were presented during this session. On day 3, participants completed extinction recall. This session was identical to the fear acquisition session except no shocks were delivered.

### Fear extinction recall analysis

Eyeblink electromyogram (EMG) responses were recorded and used to index startle responses. Initial data reduction involved averaging responses to CS+ and CS-trials within each session into blocks of two trials each. Blank screen trials within a session were averaged to acquire a baseline startle response. This baseline was then subtracted from the respective CS+ and CS-block within each session, creating scores representing potentiated startle above baseline for each CS type within each block. Note phases of the acquisition and recall sessions that contained blue or red CS+ were analyzed separately. To reduce between-subjects variability created by individual differences in startle magnitude overall, each individual participant’s blocks were standardized into Z-scores such that all scores represented departure from each individual’s mean level of potentiated startle across the entire experiment. Fear extinction recall was quantified as the initial fear response to the blue CS+ (first 4 recall session CS+ Z*-*scores) averaged together during the extinction recall session on day 3.

### Polysomnography

Polysomnography (PSG) recordings consisted of six electroencephalogram (EEG) channels (electrode positions F3, F4, C3, C4, O1, O2) referenced to contralateral mastoids (M1, M2), 2 electrooculogram (EOG) channels, and 2 submental EMG channels. Signals were recorded at 200 Hz, and subsequently exported with a 0.3-35 Hz band pass filter (plus 60 Hz notch filter) for sleep scoring. All nights of sleep were scored in accordance with American Academy of Sleep Medicine guidelines ^32^.

### Power spectral density

First, artifacts in the PSG were removed with an automated algorithm. Using frontal and central channels we calculated per-epoch summary metrics of three Hjorth parameters (signal activity, mobility, and complexity; Hjorth, 1970). Epochs where at least 1 channel was > 3 standard deviations from the mean on any of the three Hjorth parameters were marked as artifacts and removed from subsequent analysis ^33^. Artifact detection was performed over two iterations, and performed separately for each sleep stage. Power spectral density (PSD) was estimated at central electrodes (C3, C4) for all artifact free sleep separately for NREM (N2+N3) and REM, using Welch’s method with 5s windows and 50% overlap. Estimates were obtained from the temporal derivative of the EEG time series to minimize 1/f scaling. ^34^.

Given that signal amplitude is at least partly driven by individual difference factors such as skull thickness and gyral folding ^35^, we then normalized, within participant, each electrode’s power spectrum by dividing power at each frequency by that electrode’s average power (Cunningham et al., 2021; Denis et al., 2021). PSD estimates from the two central channels were averaged together to get an average estimate of spectral power.

### Statistical analysis

Statistical analysis of PSD estimates were primarily performed using cluster-based permutation testing, implemented in the FieldTrip toolbox for MATLAB ^37^. This approach allowed us to take the full power spectrum into account (and thus preserve its continuous nature), while simultaneously controlling for multiple comparisons. In other words, we were able to test our predictions about both theta and beta power within a single analysis, controlling for multiple comparisons across frequencies. Such an analysis also allowed us to detect effects in other frequency bands for which we had no *a priori* hypotheses, again without inflating the number of comparisons.

To test our primary research question, we assessed the presence of correlations between 0-30Hz PSD during sleep on the *pre-extinction* night (hereafter referred to as Night 1; **Figure 1A**) and fear extinction recall on Day 3. This analysis included participants from the Normal sleep and Post-extinction sleep deprivation groups (**Figure 1A**). The analysis was performed using the *ft_statfun_correlationT* FieldTrip function with the following parameters: 10,000 interactions, a *clusteralpha* of .05 with the default *maxsum* method to determine cluster significance, and a significance threshold of .05. Separate tests were performed for REM and NREM sleep. To better visualize significant correlations, scatterplots were also created. For these, PSD at each frequency that formed part of a significant cluster was averaged together to provide a single value for the purpose of visualization in the scatterplot. We report on PSD estimates from averaged central channels, though we note that primary results were largely unchanged when using PSD estimates from averaged frontal channels.

To test our secondary research question, we ran the same correlation analysis as above, but utilizing PSD estimates during sleep on the *post-extinction* night (hereafter referred to as Night 2; **Figure 1A**). This analysis included participants from the Normal sleep and the Pre-extinction sleep deprivation groups (**Figure 1A**). An important caveat of this analysis is that participants in the Pre-extinction deprivation group were undergoing a night of recovery sleep following total sleep deprivation the night before, and therefore may exhibit significantly altered sleep microarchitecture. To test for this, we ran a between-subjects analysis comparing PSD between the Pre-extinction deprivation group and the Normal sleep group. For these analyses, we used the same cluster-based permutation approach as the correlations, but used the *ft_statfun_indepsamplesT* function. Finally, between-group differences in the magnitudes of association were compared using the Fischer r-to-z transform, and within-group differences between nights were compared using Meng’s z test.

### Results

Group differences in extinction recall in the healthy control sample have been previously reported elsewhere (Straus et al, 2017), but we re-summarize them here (**Figure 1B**). Extinction recall was significantly impaired in the Pre-extinction sleep deprivation group compared to the Normal sleep group (*t* (44) = -2.15, *p* < .04, *d* = 0.65). The Post-extinction sleep deprivation group did not differ in extinction recall compared to the Normal sleep group (*t* (44) = -0.53, ns, *d* = 0.16).

#### Night 1

We next turned to the relationship between sleep spectral power and extinction recall. We first examined sleep PSD on Night 1, i.e. sleep following fear acquisition and prior to extinction learning, and its association with subsequent recall of extinguished fear. During REM sleep, a significant correlation cluster emerged for beta band frequencies, ranging from 19.34 - 25.97 Hz (*t*_*sum*_ = 92.63, *p* = .014; **Figure 2A**). This translated into a cluster-averaged correlation *r* = .41, *p* = .004 (**Figure 2B**), indicating that greater beta PSD during REM sleep was associated with worse recall of extinguished fear. The magnitude of the correlation was not significantly different between the Normal sleep group (*r* = .35) and the Post-extinction sleep deprivation group (*r* = .48), *z* = 0.51, *p* = .61. No other significant clusters emerged, indicating there was no association between theta band frequencies and fear extinction retention.

**Figure 2.**
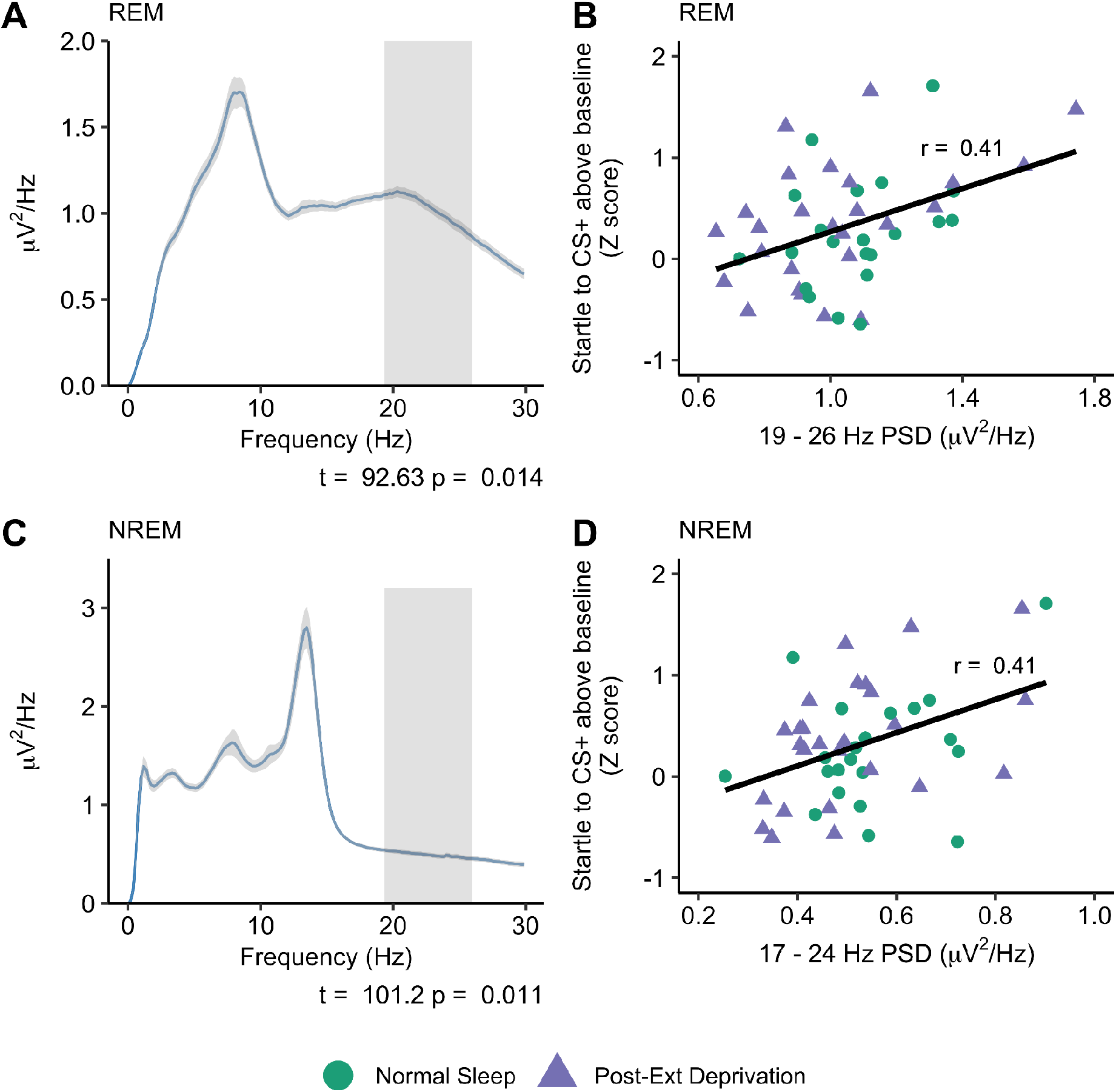
Spectral power during pre-extinction learning sleep. **A** - REM power spectrum. Frequencies showing a significant correlation with extinction recall (cluster-corrected) are highlighted in gray. Shaded area around the line indicates the between-subject standard error. **B** - Scatterplot illustrating the relationship between cluster-averaged REM PSD and fear extinction recall. **C** - NREM power spectrum. Frequencies showing a significant correlation with extinction recall (cluster-corrected) are highlighted in gray. Shaded area around the line indicates the between-subject standard error. **D** - Scatterplot illustrating the relationship between cluster-averaged NREM PSD and fear extinction recall.

With regards to NREM sleep, a near identical pattern emerged. A significant correlation cluster was found for frequencies in the beta band (16.99 - 23.63 Hz; *t*_*sum*_ = 101.19, *p* = .010), indicating that greater beta PSD during NREM sleep was also associated with worse recall of extinguished fear (**Figure 2C**). The cluster-averaged correlation magnitude was identical to REM sleep (*r* = .41, *p* = .004; **Figure 2D**), with no differences in correlation magnitude between the Normal sleep (*r* = .35) and Post-extinction sleep deprivation group (*r* = .48), *z* = 0.51, *p =* .61).

#### Night 2

We next examined PSD on Night 2, i.e. sleep following extinction learning and prior to extinction recall. No significant clusters emerged for either REM or NREM sleep, suggesting no relationship between sleep spectral power on Night 2 and extinction retention. This could be because post-extinction learning sleep is not predictive of extinction recall, or could be because one of the groups in this analysis was recovering from a night of sleep deprivation. A comparison of Night 2 spectral power between the two groups yielded no significant clusters for either REM (**Figure 3A**) or NREM (**Figure 3B**).

**Figure 3.**
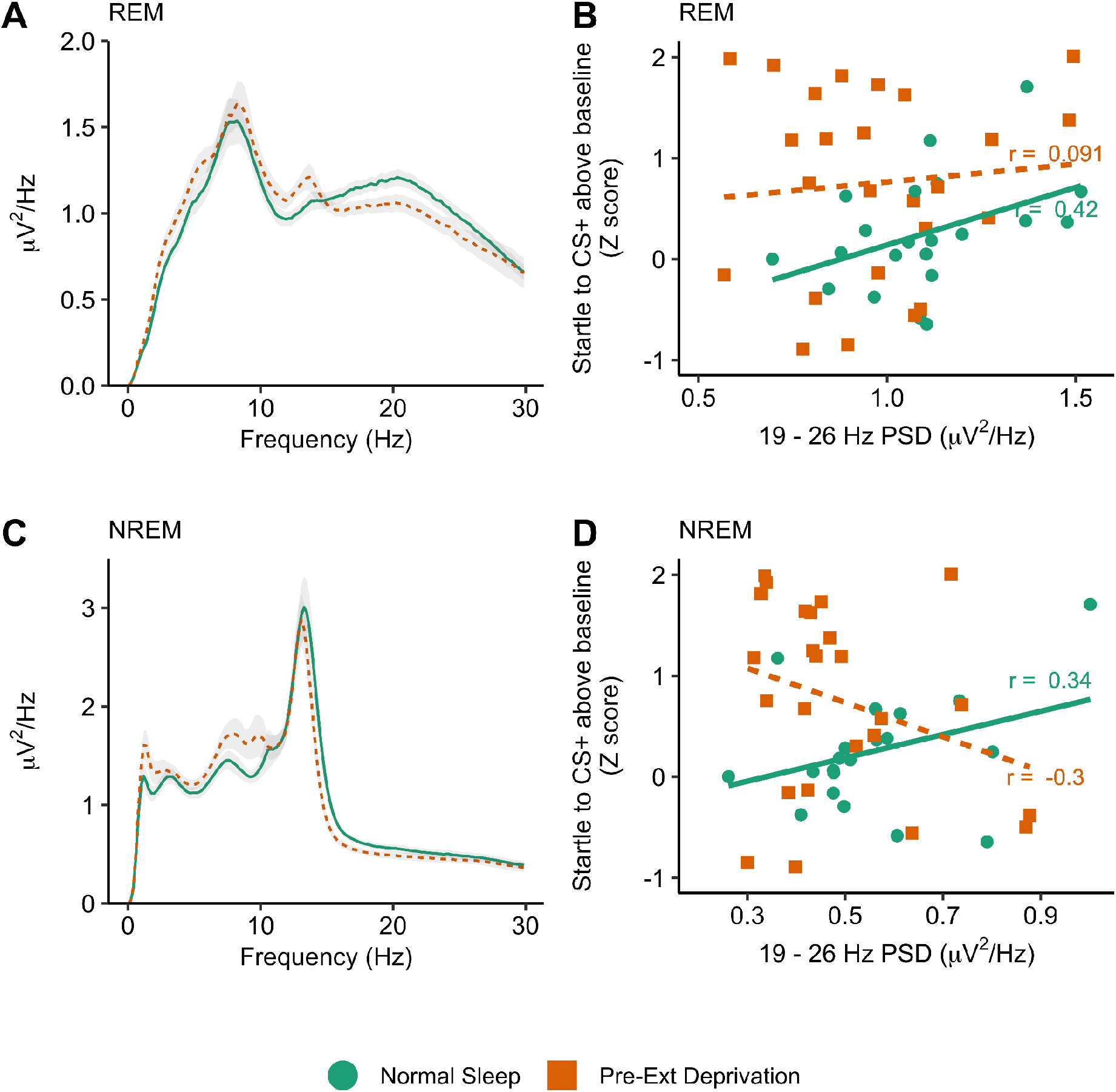
Beta power during post-extinction learning sleep. **A** - REM power spectrum. Note that the Pre-Ext Deprivation group was recovering from a night of sleep deprivation. There were no significant group differences. Shaded area around the line indicates the between-subject standard error **B** - Scatterplot illustrating the relationship between Night 2 beta PSD and fear extinction recall (frequencies derived from the significant Night 1 cluster, see Figure 2). **C** - NREM power spectrum. There were no significant group differences. Shaded area around the line indicates the between-subject standard error. **D** - Scatterplot illustrating the relationship between Night 2 NREM beta PSD and fear extinction recall (frequencies derived from the significant Night 1 cluster), see Figure 2.

When we restricted analysis to just the Normal sleep group, the correlation between beta-band PSD (collapsing across frequencies which were significant on Night 1) on Night 2 extinction recall was of similar magnitude to Night 1 (REM: *r* = .42; NREM: *r* = .34). For both sleep stages, the difference in correlation magnitude was not significant (REM: *z* = 1.02, *p* = .31; NREM: *z* = 0.13, *p* = .90).

In the Pre-extinction deprivation group, the correlation between Night 2 beta power and extinction recall was not statistically significant, showing either no (REM: *r* = .09, *p* = .67; **Figure 3C**) or a non-significant negative relationship (NREM: *r* = -.30, *p* = .15; **Figure 3D**). For NREM sleep, there was a significant difference in correlation magnitude between the two groups (*z* = 2.09, *p* = .037). The REM sleep correlation magnitudes were not significantly different from each other (*z* = 1.02, *p* = .31)

#### Stability of beta PSD across nights

Finally, we examined whether beta PSD estimates were stable across nights. Using the Normal sleep group, we were able to investigate the stability of sleep beta activity across the three experimental nights (Baseline, Night 1, Night 2; both sleep deprivation groups were excluded from this analysis). There was no significant difference in beta power between the three nights for either REM (*F* (2, 40) = 1.88, *p* = .166, *np*^*2*^ = .086) or NREM (*F* (2, 40) = 0.68, *p* = .512, *np*^*2*^ = .033) sleep (**Figure 4A-B**). Next, intraclass correlation coefficients (ICC) were used to quantify the level of similarity of an individual’s beta power across the three nights. ICCs were found to be very high (REM ICC = 0.94 [0.89, 0.97]; NREM ICC = 0.93 [95% CI: 0.87, 0.97]), indicating a high agreement in beta PSD across nights (see **Figure 4C-D** for illustrative examples).

**Figure 4.**
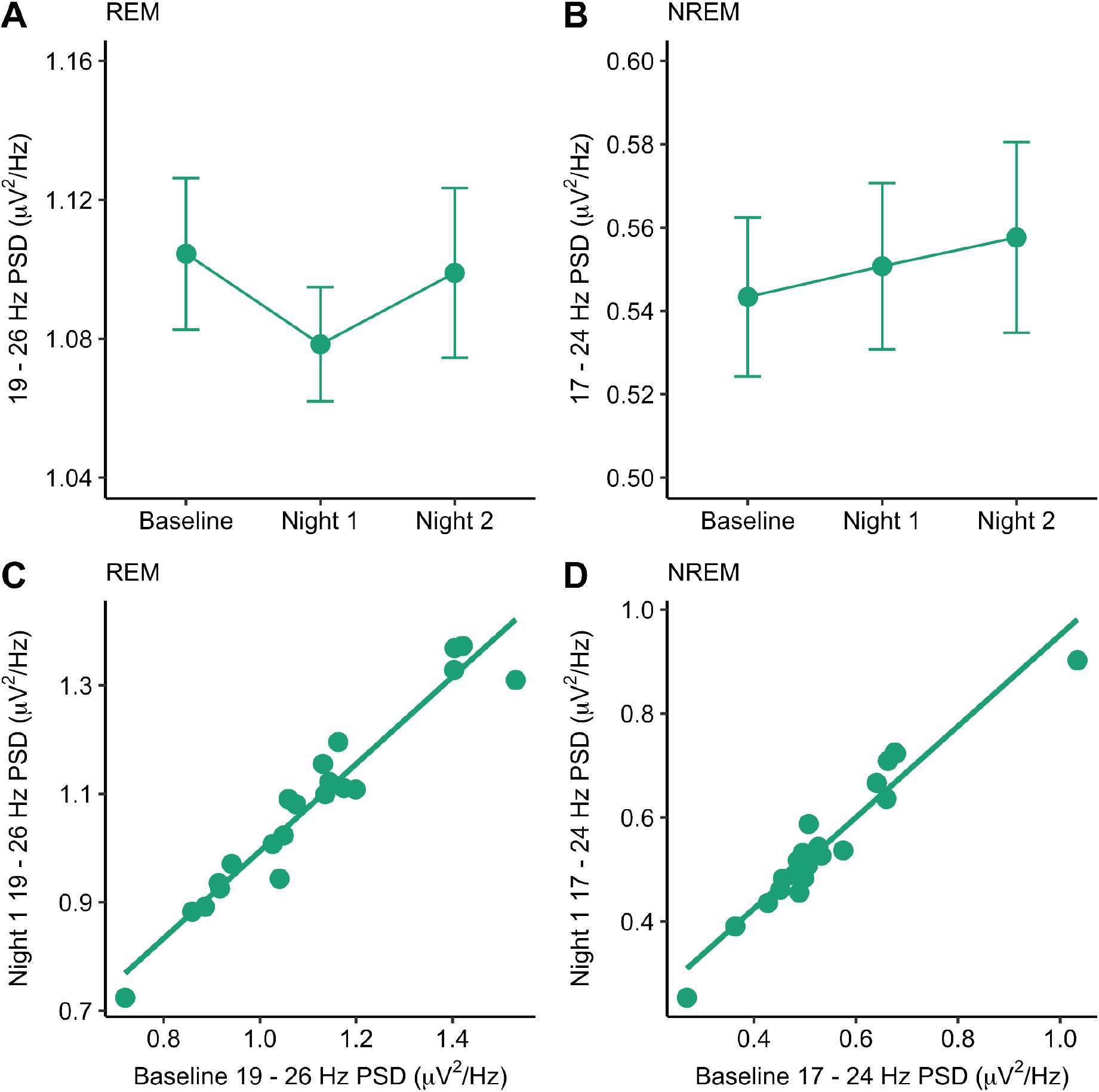
Beta PSD during sleep is stable across nights. **A** - Beta power during REM sleep across three nights in the normal sleep group. **B** - Beta power during NREM sleep across three nights in the normal sleep group. **C** - Correlation between REM beta power during the baseline night and during night 1 (pre-extinction learning). **D** - Correlation between NREM beta power during the baseline night and during night 1 (pre-extinction learning). Error bars indicate the within-subject standard error.

### Interim summary

These results suggest that in healthy participants, heightened beta power during both NREM and REM sleep were associated with impaired recall of extinguished fear. As such, high beta power during sleep exert a similar negative effect on extinction recall as a night of sleep deprivation. Beta PSD estimates were highly consistent across nights, suggesting them to be a stable marker of an individual’s sleep microarchitecture.

### Study 2 Methods

This study was a reanalysis of a previously published dataset ^29^ of individuals with PTSD. Please see Straus et al (2018) for full details. The investigation was carried out in accordance with the Declaration of Helsinki. The VA Internal Review Board as well as the University of California, San Diego’s Human Research Protections Program approved the study. Informed consent was obtained from all participants.

### Participants

Study 2 participants were 15 Veterans with a primary diagnosis of PTSD. A total of 12 participants with complete, usable data were included. Participants had no history of psychosis, substance use disorders in the six months prior to the study, and no untreated sleep disorders other than insomnia and nightmares. Included participants exhibited a consistent startle response at screening (over 75% discernable responses to twelve 105dB 40ms startle pulses).

### Procedure

Participants followed the same procedure as the Normal sleep group of Study 1 (**Figure 1A**), except the baseline night also served as an adaptation night.

### Fear potentiated startle

Fear acquisition started with the same six trial acclimation period as Study 1. Following acclimation, participants were exposed to: 1) eight 6-second long presentations of a blue circle (CS+), paired with an air puff (US) to the throat in 75% of trials; 2) eight 6-second long presentations of a yellow circle that was never paired with an air puff (CS-); and 3) 8 presentations of the startle pulse in the absence of any stimuli. Stimulus presentation was block randomized with the constraint of two trials of each type per block. On day 2, participants underwent extinction learning, consisting of 16 presentations each of CS+, CS-, blank screen along with startle pulses but without any air puffs being delivered. On day 3, participants performed the extinction recall session. This session followed the exact same procedure as initial fear acquisition, except no air puffs were delivered.

### Statistical analysis

Analysis of extinction recall and spectral analysis were identical to Study 1. To test for group differences, we compared Night 1 spectral power between Healthy controls (Normal sleep and Post-extinction deprivation groups only) and individuals with PTSD, using the same cluster-based permutation framework as in study 1, with the *ft_statfun_indepsamplesT* function to compare the two groups. We focused our group difference analysis on Night 1 rather than the Baseline night because Baseline night also served as adaptation night for the PTSD group (**Figure 1A**), meaning that any group differences could have been due to a first-night effect. Equally, by performing our group difference analysis on Night 2, our healthy control sample would have contained participants who were recovering from a night of sleep deprivation, and removing this group would greatly reduce our statistical power. To obtain an effect size for each significant cluster, PSD values for each frequency within a given cluster were averaged together, and the HC vs PTSD group difference was compared with Cohen’s *d*.

We examined the relationship between sleep spectral power and extinction recall in PTSD by correlating PSD averaged across frequencies which revealed a significant group difference in the above analysis. Because our analysis of healthy controls found associations with beta band power only, in order to limit the number of comparisons, we focused our correlational analysis only on group-differences that emerged in the canonical beta band (∼16 - 30Hz).

## Results

### Group differences

During REM sleep, we found significantly higher PSD in the beta frequency range (17.58-23.83Hz; t_sum_ = 105.15, *p* = .007, *d* = 0.90; **Figure 5A**) in PTSD relative to controls. We additionally observed significantly lower high theta/low alpha REM PSD in PTSD (7.03-10.94Hz; t_sum_ = 68.03, *p* = .030, d = 1.22). When we assessed group differences in NREM sleep, we again found significantly higher beta PSD in PTSD relative to controls (15.82-21.48Hz; t_sum_ = 81.13, *p* = .038, *d* = 0.62; **Figure 5B**).

**Figure 5.**
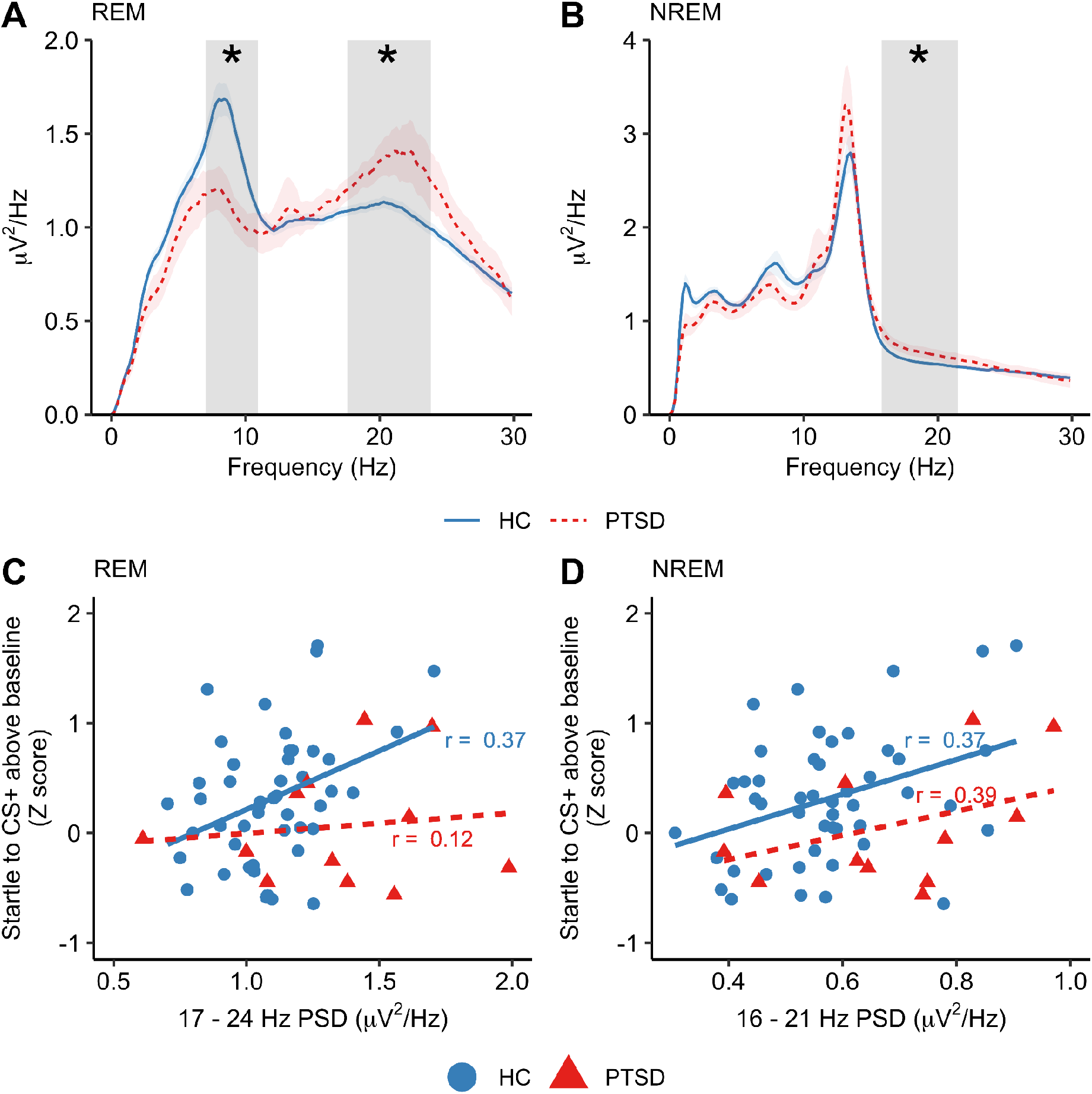
Spectral power differences in PTSD. **A** - REM power spectrum. Significant group differences between healthy controls and PTSD highlighted in gray. Shaded areas around the line indicate the between-subject standard error. **B** - NREM power spectrum. Significant group differences between healthy controls and PTSD highlighted in gray. Shaded areas around the line indicate the between-subject standard error. **C** - Correlation between REM beta and extinction recall in healthy controls (solid blue line) and PTSD (dashed red line). **D** - Correlation between NREM beta and extinction recall in healthy controls and PTSD. * = *p* < .05, + = *p* < .10.

### Correlations with extinction recall

We correlated cluster-averaged Night 1 beta power (i.e. beta-band frequencies which significantly differed between the groups) with fear extinction recall. We did not observe a correlation between beta power during REM sleep and extinction recall in the PTSD sample (*r* = .12 *p* = .704; Healthy controls: *r* = .37, *p* = .011; **Figure 5C**). For the NREM cluster however, a positive correlation with a magnitude similar to the healthy controls was observed (PTSD: *r* = .39, *p* = .214; Healthy controls: *r* = .37, *p* = .011; **Figure 5D**). The lack of statistical significance in the PTSD group may be attributed to the small sample size. Similar patterns in the PTSD group emerged when we ran the analysis on Night 2 PSD estimates (REM: *r* = .28, *p* = .40; NREM: *r* = .42, *p* = .20).

### Stability of beta PSD estimates in PTSD

Finally, we examined whether PSD estimates in the frequency band associated with impaired extinction recall is also a stable measure in PTSD. As with healthy controls, beta spectral power was found to be highly stable across all three nights in PTSD (REM: ICC = .94 [.87, .97]; NREM: ICC = .92 [95%CI: .78, .98]). In summary, this provides the first, albeit highly preliminary, evidence that beta frequency activity during sleep, characteristic of PTSD sleep, is indeed associated with the often-documented deficit in fear extinction recall seen in this group.

## Discussion

We found that increased beta spectral power during pre-extinction learning sleep (Night 1) was associated with poorer extinction recall. This association was significant for both NREM and REM sleep. This mirrors that of prior work using sleep deprivation, which found an impairment in extinction recall when total sleep deprivation preceded, rather than followed, initial extinction learning ^15^. Pre-encoding sleep loss has been linked to attenuated functioning in memory-related brain structures, such as the hippocampus ^38–40^, as well as impaired consolidation, even following subsequent periods of recovery sleep ^40^. In a fear conditioning task, neuroimaging work has shown a failure to activate both fear and extinction processing-related brain regions following either sleep restriction or total sleep deprivation ^41^. Similarly, beta power during sleep has also been suggested as a proxy of disturbed sleep, and is related to increased autonomic arousal and the disturbance of both NREM and REM sleep ^30^. As such, it is possible that sleep disturbance, indexed here as heightened beta spectral power, led to changes in the neural substrates of extinction learning, which in turn led to poor consolidation of the fear extinction process.

A notable finding was that Night 1 beta spectral power predicted extinction recall in the post-extinction deprivation group, who went on to be sleep deprived the following night (Night 2). In the Normal sleep group, the relationship between beta spectral power and extinction retention was equivalent in magnitude on both Night 1 and Night 2, suggesting that beta spectral power levels consistently predict extinction recall ability, independently of when sleep occurs relative to extinction learning. This result is consistent with our finding that beta PSD estimates were highly consistent between nights.

The high night-to-night consistency of beta PSD in both healthy controls and PTSD aligns with a large body of work showing the sleep EEG to be a highly stable and heritable trait ^33,42–44^. Higher beta power during sleep (relative to healthy controls) has been documented in a number of psychopathologies, including PTSD ^18,19,21^. Notably, when individuals with PTSD are compared to trauma-exposed, non-PTSD participants, higher beta power is not always observed ^27,28,45^. Prospective, longitudinal research in this area may be able to identify certain sleep oscillatory activity as a characteristic biomarker with regards to the risk or resilience toward the onset of PTSD following a trauma exposure.

As well as finding beta spectral power to be heightened in PTSD relative to healthy controls, we also found that the association between pre-extinction learning NREM beta power was associated with poorer extinction memory at a magnitude similar to healthy controls. We therefore speculate that beta spectral power may be a stable sleep biomarker that confers vulnerability to impaired consolidation of fear extinction. While this result is both novel and intriguing, we stress that these results should be considered preliminary due to the small sample size, and so must be replicated in larger samples. While the relationship between NREM beta power and extinction recall showed a similar strength of association in both healthy participants and PTSD, this was not the case for REM sleep, where the correlation was smaller in PTSD. This could reflect a functional dissociation in NREM vs REM sleep microarchitecture following a trauma, especially given that numerous alterations in REM architecture and physiology have been observed in PTSD ^4^. Furthermore, work in other clinical groups such as insomnia has observed opposing relationships between REM sleep and fear extinction recall between patients and controls ^8^.

We found novel evidence that beta band spectral power during sleep impairs fear extinction recall. However, the mechanistic basis and origins of the beta rhythm in the sleep EEG remains poorly understood. Beta activity is often cited as a marker of cortical hyperarousal, and has frequently been shown to be heightened in insomnia ^46^. Such a link is not always found however. For example, following a mindfulness-based intervention for chronic insomnia, NREM beta power increased from baseline and was associated with fewer symptoms ^47^. In a large sample of PTSD patients and trauma-exposed controls, beta power levels were associated with fewer symptoms of subjective hyperarousal, fewer nightmares, and improved emotion regulation ^27^. As such, it has been proposed that while beta activity during sleep may index heightened arousal, the nature of this hyperarousal may either be adaptive or maladaptive ^47^.

While we found that beta band activity was associated with impaired extinction recall, we did not find any evidence of an association between REM theta (∼4-8Hz) PSD and extinction recall. Some studies have found a link between emotional memory recollection and REM theta power ^48,49^, especially when emotional memories are encoded under stressful conditions ^50^. However, this link is not consistent across the existing literature ^26^, and enhancing REM theta oscillations through acoustic stimulation does not lead to expected improvements in emotional memory ^51^. Some work has also suggested that the magnitude of REM theta oscillations may be predictive of resilience to developing PTSD after trauma exposure. The current data however do not support a role of REM theta power in extinction retention processes. As such, future work should explore other mechanisms by which REM theta may be protective against PTSD development.

The present study has several limitations that must be acknowledged. First, there were slight differences in the fear conditioning protocol between Study 1 (Healthy controls) and Study 2 (PTSD). In Study 1, the unconditioned stimulus consisted of an electric shock, whereas in Study 2 consisted of an air puff. Although both approaches are widely used in the fear conditioning literature, this methodological difference may account for why, at a behavioral level, extinction recall was no worse in PTSD compared to controls. Although this difference is a limitation, we note that our primary goal here was not to compare group differences in extinction recall itself, but rather determine associations between sleep spectral power and the ability to recall extinguished fear. The fact that, for NREM sleep at least, correlations were of a similar magnitude in both groups suggests similar sleep mechanisms contributed to extinction recall processes in the two studies. Future work, using primary datasets where methodologies are equivalent across all groups, should aim to replicate these findings.

Another limitation is that the association between beta power and extinction recall is purely correlational, and as such causal inferences cannot be conclusively made from this data alone. Acoustic and electrical stimulation methods can be used to enhance frequency-specific sleep oscillations that have been shown to either promote ^52^ or impair ^53^ memory consolidation, and are a fruitful target for future research. A final limitation is that we did not collect any measures of insomnia symptom severity, meaning we were unable to investigate whether beta spectral power was linked to any symptoms of hyperarousal seen in insomnia.

## Conclusion

Sleep disturbances are extremely common in PTSD, and contribute to its onset and maintenance. We found evidence that beta spectral power was associated with subsequent ability to recall extinguished fear in healthy controls and individuals with PTSD. Given the importance of extinction recall for the treatment of PTSD symptoms, this work provides an important new contribution to our understanding of the electrophysiological underpinnings of a key mechanism of PTSD maintenance.

## Acknowledgements

This research received support from the Department of Defense Grant No. DM102425 (to SPAD) and National Institute of Mental Health National Research Service Award No.

F31MH106209 (to LDS) and the Veterans Affairs Center of Excellence for Stress and Mental Health.

